# A Minimal Packaging Signal Enables Production of High-Purity Phage-Like Particles for CRISPR-Cas Antimicrobials in *Staphylococcus aureus*

**DOI:** 10.64898/2026.06.26.734846

**Authors:** David S. Dooley, Hannah Boyd, Cong T. Trinh

## Abstract

Precision phage therapeutics provide a promising strategy to combat multidrug-resistant pathogens, including *Staphylococcus aureus*. Efficient, specific packaging of genetic cargoes remains challenging. Using modular design principles, we report a minimal phagemid packaging signal consisting of the phage terminase small subunit under its native promoter that significantly outperforms conventional packaging signals. The utility of this synthetic *terS* operon was demonstrated through production of highly concentrated and genetically pure CRISPR-Cas antimicrobials. To circumvent CRISPR-mediated self-targeting during antimicrobial generation, a *terS*-deficient strain was engineered to express the anti-CRISPR protein AcrIIA4, enabling titers above 10^10^ transducing units per milliliter (TRU/mL) with over 94% purity. With a high-copy origin of replication module, CRISPR-Cas phage-like particle titers could approach 10^12^ TRU/mL. We discovered that pure CRISPR-Cas antimicrobials are potent and can be amplified in hosts possessing prophages. Taken altogether, this study defines the minimal and optimal genetic requirements for efficient, specific creation of phage-based technologies.

**Technological Readiness:** The described system for engineering phage-like particles has reached a technological readiness level (TRL) of 4–5 based on the provided laboratory validation and strong literature support from other engineered phage therapies applied to *in vivo* models. Previous systems have demonstrated high-purity phage-like particle preparation, but always at the cost of severely reduced productivity. Therefore, our demarcation of a specific, efficient, and minimal system for packaging nucleic acid cargoes into phage vectors is a critical step toward real-world use. Despite this, low levels of contaminating host/phage DNA remain a key barrier to phage-based therapies. Protein and strain engineering efforts can help mitigate terminase nonspecificity, but care must be taken to not compromise productivity. More generally, widespread adoption will require deeper understanding of host-pathogen-phage interactions, development of scalable GMP manufacturing processes, and harmonized regulatory guidance that recognizes the dynamic nature of phage-derived technologies.

## Introduction

Originally identified in 1915[1] and first used to treat bacterial infections in 1919[2], bacteriophages have long been appreciated as highly specific and lethal predators of bacteria. Recent advances in synthetic and systems biology have allowed unprecedented insights into their intricate physiology, opening doors to expanded use in agricultural[3], environmental[4], and medical[5] sectors. To increase their utility, phages may be engineered with genetic payloads that can degrade biofilms[6], inhibit cell-to-cell communications[7], resensitize antibiotics[8], or improve lethality[8, 9]. Typically, these payloads are incorporated directly into the phage genome, but editing lytic phages can be onerous and phage genomes frequently possess undesirable toxins and virulence genes[10, 11]. Therefore, understanding and controlling the efficiency and specificity of packaging payloads is essential for precision therapy.

One promising strategy to generate high-purity phage treatments is to encode therapeutic components, such as CRISPR-Cas antimicrobials[8, 12–15], on a phagemid for packaging into the phage’s capsid. This design strategy has the potential to produce targeted therapies that are free of unwanted host or phage genomes, but gaps in understanding limit this approach. For example, it is not well characterized which genetic elements must be included on the phagemid to enable highly efficient and specific packaging of therapeutic cargoes into phage-like particles (PLPs). It also remains unknown how recombinant hosts need to be modified to avoid nonspecific packaging.

In the human pathogen *Staphylococcus aureus* and its siphoviruses, such as *ϕ*NM1 of the Newman strain, the most common packaging signal is the *rinA*-*terS*-*terL* operon that encodes a phage transcriptional activator (RinA) and the terminase complex (TerS-TerL) responsible for recognizing and loading procapsids with phage DNA[8]. Some have suggested that inclusion of an additional gene, *rinB*, is necessary for optimal phagemid packaging, although its exact role in packaging and extent of its benefit remain in question[14].

Beyond packaging efficiency and specificity, controlling the size of packaging signals is critical but underappreciated. For instance, inclusion of *rinA* and *rinB* increases the size of the packaging signal by ∼1 kilobase (kb), bringing its total size to ∼2.7 kb. Once other essential elements have been included, like resistance marker(s), origin(s) of replication, and therapeutic payload(s), phagemid size can become exceedingly large. Because increasing phagemid size complicates cloning efforts and compromises packaging efficiency and fidelity[8, 14], it is critical to determine the phagemid packaging sequence(s) that are both minimal and optimal.

Phagemid packaging signals must be paired with genetic manipulation of production strains to prevent unwanted incorporation of phage genomes in treatments. Because the *pac* site that is recognized by the terminase complex is embedded within *terS*[16, 17], deletion of the prophage *terS* from the production strain is critical for generating pure PLPs. If the *S. aureus* production host does not contain the deletion of *terS*, it has been shown that up to 97% of PLPs can package wild-type (WT) phage genomes, diluting treatments and opening the possibility for virulence factor transfer and toxin expression[8].

In this study, using *S. aureus* and its siphoviruses as a model system, we define a minimal packaging signal required for efficient and specific phagemid packaging. We accomplish this by performing systematic characterization of a modular phagemid packaging signal in both wild-type and Δ*terS* production strains. This minimal and optimal packaging signal is a modular synthetic operon comprising *terS* and its cognate promoter. Using this system, we demonstrate high-purity and high-titer production of phage-like particles harboring CRISPR-Cas antimicrobials. To avoid self-targeting, we engineer an anti-CRISPR-harboring Δ*terS* production strain to produce these CRISPR-Cas antimicrobials. We further demonstrate that development of a high-copy phagemid can greatly improve PLP titer, which allows for higher dosing to overcome limitations in pure CRISPR-Cas antimicrobials. We found that PLPs harboring CRISPR-Cas antimicrobials can self-replicate when treating *S. aureus* pathogens with resident prophages. We envision that this study establishes a foundation for high-purity, high-efficiency phage-based delivery of nucleic acid cargoes, with applications in precision phage therapy and antimicrobials.

## Results and Discussion

### Define a minimal packaging signal module to produce pure phage-like particles in *S. aureus*

#### Design of a minimal packaging signal module and production host

The phagemid packaging signal is required to contain a *pac* site, which is embedded in the *terS* gene for packaging by *ϕ*NM1 of the Newman strain[16, 17]. We designed phagemids with a modular packaging signal consisting of a *terS* operon with either i) no promoter, ii) a strong constitutive promoter (P*_rpsL_*), or iii) 86 bp of its native promoter (P*_terS_*) (**Fig. 1A**). As controls, we included an empty vector and *terS-terL* operons under the same expression conditions as *terS*. To determine a minimal phagemid packaging signal in *ϕ*NM1, we also constructed a *terS*-deficient strain by deleting *terS* from a single *ϕ*NM1 lysogen strain (SaDD0001) derived from the prophage-free, restriction-free *S. aureus* RN4220[18], using pCasSA with 500 base pair (bp) homology arms[3] (**Fig. S1**). The resultant strain, *S. aureus* SaDD0003, was confirmed via PCR, sequencing, and phenotypic screening (**Figs. 1B–1C**). The phagemids were transformed into either *S. aureus* SaDD0001 or SaDD0003, confirmed by colony PCR, and used to generate PLPs. PLPs were titered for both transducing units (TRU) and plaque-forming units (PFU) on SaDD0001 and RN4220, respectively. Transduction quantifies the packaging efficiency of phagemids based on the transfer of an antibiotic selection marker while plaque formation measures natural phage genomes capable of causing infection events.

**Figure 1:**
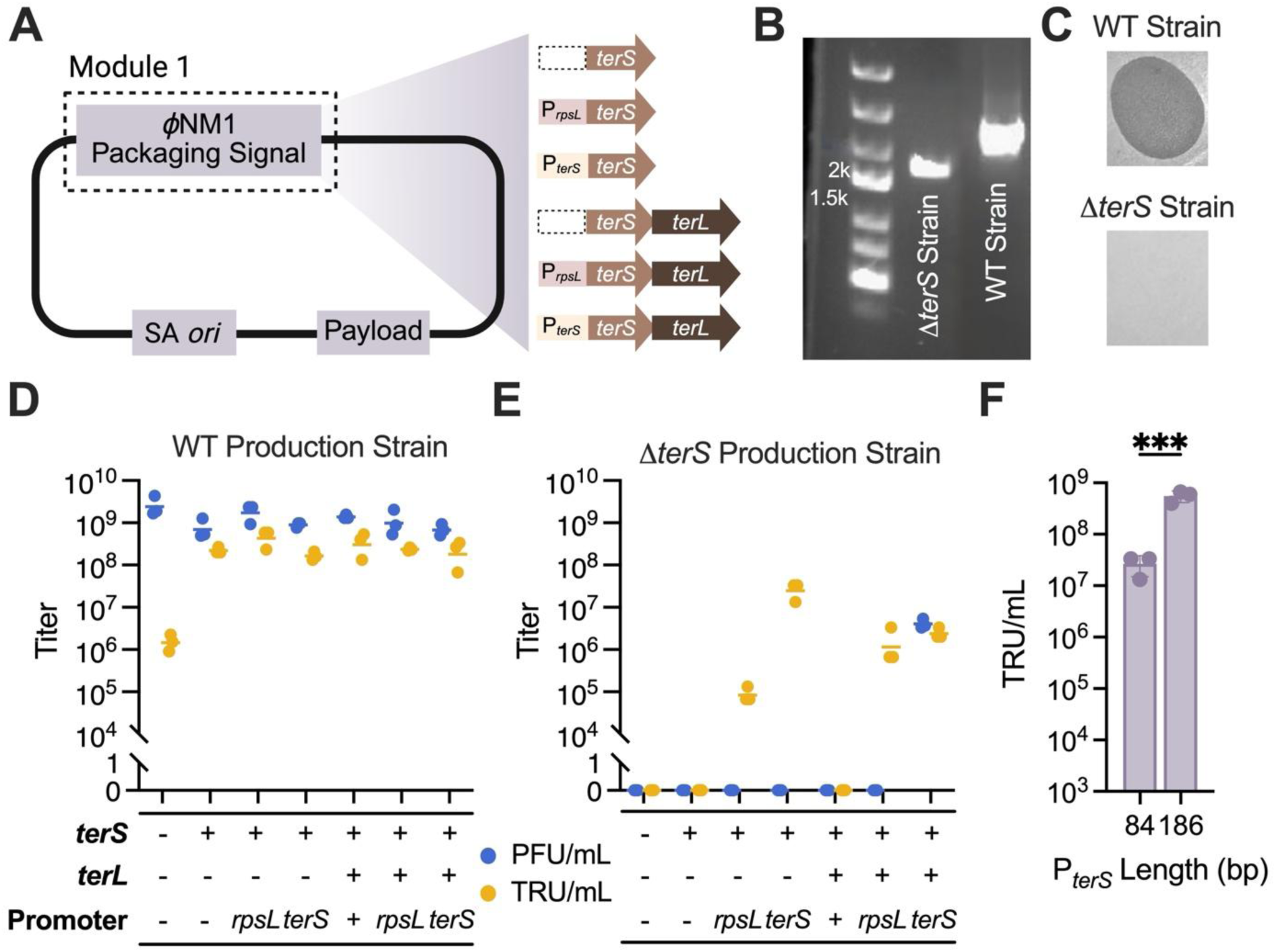
*TerS* under its native promoter is sufficient for phagemid packaging in a Δ*terS* production strain SaDD0003. (**A**) Modular phagemid design for *ϕ*NM1 packaging signal. (**B**) DNA gel of PCRs confirming *terS* deletion. (**C**) Plaque-forming phenotypes of lysates generated from the parent strain SaDD0001 and Δ*terS* strain SaDD0003. (**D–E**) Titer of plaque-forming units (blue circles) and transductants (yellow circles) per milliliter of phage lysate generated from either (**D**) parent or (**E**) Δ*terS* production strains harboring phagemids with various packaging signals. (**F**) Comparison of truncated and full-length *terS* promoters on packaging efficiency. Statistical significance determined by an unpaired t-test assuming a lognormal data distribution.

#### Phagemid packaging is promiscuous by wild-type phage, and terS is essential for specific packaging of nucleic acid cargoes

To determine the promiscuity of phagemid packaging by the wild-type phage *ϕ*NM1, we induced SaDD0001 harboring an empty vector with mitomycin C for the production of phages and PLPs. SaDD0001 was able to produce an average of 2.63 ± 1.47×10^9^ PFU/mL (**Fig. 1D**). Low-level generalized transduction of the empty vector was observed due to promiscuous packaging, with a mean TRU/mL of 1.56 ± 0.67×10^6^. A phagemid with *terS* but no promoter increased transduction efficiency by 2.15 logs to 2.22 ± 0.39×10^8^ TRU/mL, consistent with the phage *pac* site being embedded within *terS*[16, 17]. Additional expression of *terS* from the phagemid by either P*_rpsL_* or P*_terS_* did not increase phagemid packaging efficiency, nor did the incorporation of *terL*, indicating that sufficient terminase proteins are produced from the prophage (**Fig. 1D**). It is also noted that phagemids with *terS* alone had the highest ratio of phagemids to WT phage genomes (TRU/PFU) at 35.5%, which was higher than reported for a packaging signal of *rinA*-*terS*-*terL* in a phagemid of similar size (24%)[8]. Taken altogether, these results indicate that the wild-type phage *ϕ*NM1 randomly packages cargoes at low levels, and the *terS* sequence is essential for efficient cargo packaging.

#### Characterization of a minimal packaging signal in S. aureus

To minimize nonspecific packaging, we characterized the modular packaging signals in SaDD0003. As a negative control, phage lysates induced from *S. aureus* SaDD0003 harboring an empty vector did not form plaques or transductants, underscoring the necessity of the TerS protein for DNA packaging into phage particles (**Fig. 1E**). Similarly, complementation of *terS* without a promoter was unable to restore DNA packaging. As expected, transduction—but not plaque formation—was rescued with the addition of P*_rpsL_* before *terS*, albeit at a lower level than nonspecific phagemid packaging in SaDD0001 (**Fig. 1D, 1E**), indicating the level of *terS* expression is critical for efficient DNA encapsidation. We found that when *terS* expression was driven by the native promoter, P*_terS_*, transducing titers were 2.48 logs higher compared to P*_rpsL_* while still excluding WT phage genomes (**Fig. 1E**). On average, P*_terS_*-*terS* lysates from SaDD0003 contained 2.66 ± 1.15×10^7^ TRU/mL, compared to 1.67 ± 0.37×10^8^ TRU/mL for lysates generated from SaDD0001, a 0.80 log_10_-fold decrease (**Figs. 1D, 1E**). We hypothesized that this difference was likely due to either modified expression of *terL* in SaDD0003, or more likely, partial activation by RinA since 102 bp between *rinA* and *terS* were not included in our original P*_terS_* design. In support of the latter hypothesis, the full 186-bp *terS* promoter region was cloned in front of *terS* and assayed for packaging efficiency, revealing a mean TRU/mL of 5.56 ± 1.39×10^8^, which was 1.32 logs higher than the 86-bp P*_terS_* and equivalent to the packaging efficiencies observed for *terS* phagemids in SaDD0001 (**Fig. 1F**).

To further demonstrate that the synthetic operon of *terS* is minimal for specific packaging, we analyzed the effect of introducing *terS-terL* without and with various promoters. Addition of *terL* after *terS* without a promoter yielded lysates without detectable TRU or PFU, and again, P*_rpsL_* reinstated transducing ability when driving *terS*-*terL* expression (**Fig. 1E**). Notably, the *terS*-*terL* packaging signal outperformed *terS* alone by 1.24 logs when expressed from P*_rpsL_*. However, P*_terS_* was unable to improve the performance of *terS*-*terL* as it did for *terS*, only increasing TRU/mL by 0.20 logs (**Fig. 1E**). When *terS-terL* was complemented under P*_terS_*, plaque-forming ability was restored at low levels (4.11 ± 1.07×10^6^ TRU/mL in SaDD0003 vs. 7.00 ± 2.18×10^8^ in SaDD0001). This is the only packaging signal with both upstream and downstream homology for *terS*, leading to homologous recombination and repair of Δ*terS* prophages. This finding corroborates previous reports showing that removal of homologous regions around the complemented phage packaging signal is necessary for preventing natural phage contamination[19].

Taken altogether, a synthetic operon containing *terS* with its full 186-bp *terS* promoter defines a minimal packaging signal in *S. aureus*. This unique design is different from the *rinA*-*terS*-*terL* operon that has been used in *S. aureus*, which is not minimal. The use of this synthetic operon greatly reduces phagemid size, which can be used to load necessary gene functions for delivery. It is critical to produce these PLPs in *terS*-deficient strains to avoid nonspecific packaging. Furthermore, eliminating *terL* in the synthetic operon avoids nonspecific packaging of the native phage genome in the *terS*-deficient strain due to homologous recombination. Previous studies have claimed that *terS* alone cannot efficiently package phagemids in temperate staphylococcal phages[14, 19]. To the best of our knowledge, however, these studies did not place *terS* under an active promoter, likely resulting in poor protein expression and packaging rates in the Δ*terS* strain (<10^6^ TRU/mL). When complemented in its native genomic context (i.e., its promoter region), *terS* enabled efficient packaging of phagemids while preventing the inclusion of contaminating phage genomes. Although *rinA* and *rinB* are likely important for efficient DNA packaging, they are not known to play a role in determining which nucleic acids are packaged; therefore, expression of these proteins from the prophage instead of the phagemid can minimize size constraints on therapeutic phagemids without hindering phagemid packaging or favoring natural phage production.

### Minimal packaging signal enables production of high-purity, high-titer CRISPR-Cas antimicrobials

We next investigated whether a minimal packaging signal can be utilized to produce CRISPR-Cas antimicrobials with high purity and efficiency that can be used to inactivate *S. aureus*. These CRISPR-Cas antimicrobials are PLPs harboring CRISPR-Cas systems that can target any locus, causing chromosomal damage and disrupting gene function, thereby inactivating the pathogens. To produce high-purity, high-titer CRISPR-Cas antimicrobials, we engineered a host strain capable of packaging therapeutic phagemids without self-targeting or wild-type phage contamination.

#### Engineering of CRISPR-Cas antimicrobials and an AcrIIA4-harboring, terS-deficient S. aureus host for producing pure CRISPR-Cas antimicrobials without self-targeting and promiscuous packaging

To demonstrate the utility of our minimal packaging signal to produce CRISPR-Cas antimicrobials, we designed a modular phagemid with a fixed minimal packaging module and variable payload module, encoding either an active or null CRISPR-Cas system (**Fig. 2A**). The 86-bp P*_terS_*-*terS* packaging signal module was used in our design described above, while the module of SpCas9 and guide RNA (gRNA) was placed under constitutive promoters to mirror systems that could be deployed in real-world treatment scenarios.

**Figure 2:**
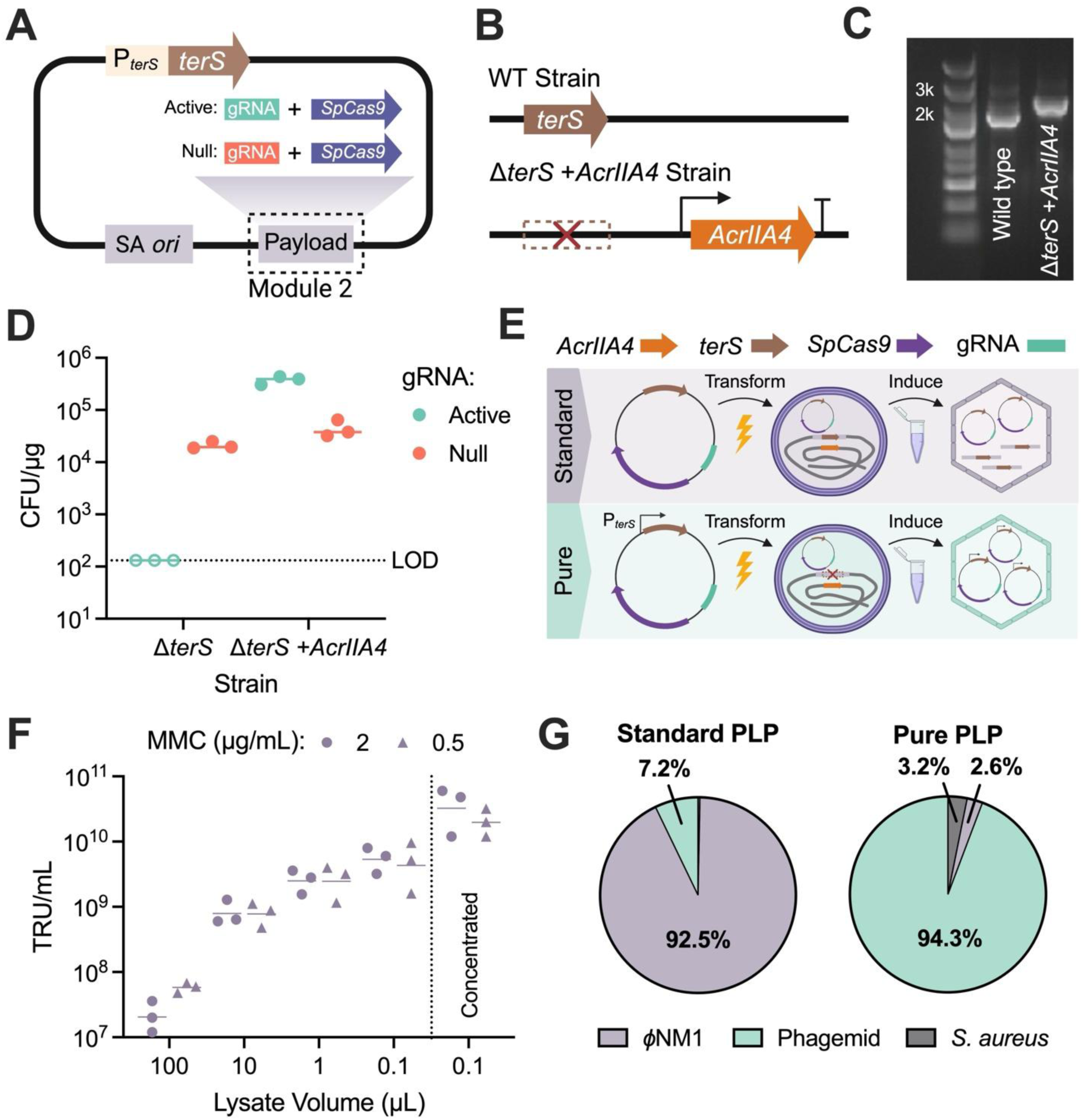
High purity and titer of effective CRISPR-Cas antimicrobials with minimal contamination can be generated with the P*_terS_*-*terS* packaging signal in the *terS*-deficient, AcrIIA4-harboring production strain SaDD0007. (**A**) Modular phagemid design for CRISPR-Cas antimicrobials. (**B**) Engineering the Δ*terS* +*AcrIIA4* production strain SaDD0007. (**C**) DNA gel of PCRs confirming *AcrIIA4* insertion into the production strain. (**D**) Transformation efficiencies of phagemids harboring either active or null-targeting gRNAs into *S. aureus* without *terS* (Δ*terS*) or *S. aureus* without *terS* and with *AcrIIA4* (Δ*terS* +*AcrIIA4*). Open circles represent measurements below the limit of detection (LOD). (**E**) Workflow demonstrating the differences between standard and pure PLP production. (**F**) Effect of inducer concentration, lysate volume, and ultracentrifugation on PLP phagemid titer. (**G**) Relative abundance of encapsidated DNA in standard and pure PLP formulations.

A corollary of fully constitutive expression is that phagemids with gRNAs targeting the host chromosome (i.e., “self-targeting” gRNAs) are highly toxic, preventing the production of therapeutic PLPs in an *S. aureus* production host. Indeed, when *S. aureus* SaDD0003 was transformed with a self-targeting CRISPR-Cas phagemid, no transformants were recovered (**Fig. 2D**). To circumvent this barrier, AcrIIA4 was integrated into the Δ*terS* production strain under P*_SarAP1_*, resulting in *S. aureus* SaDD0007 (**Figs. 2B–2C**). AcrIIA4 provided robust protection against a self-targeting phagemid, enabling a mean transformation efficiency of 3.79 ± 0.67×10^5^ CFU/µg, which surpassed even the non-targeting (NT) control (**Fig. 2D**). *S. aureus* SaDD0007 was subsequently used as a production strain for pure CRISPR-Cas antimicrobials by transforming it with the phagemids and inducing with mitomycin C (MMC).

#### Ultracentrifugation is critical to remove lysate contaminants and enhance the purity and titer of PLPs harboring CRISPR-Cas antimicrobials

Even though the design of SaDD0007 and CRISPR-Cas antimicrobials can improve specific packaging and avoid self-targeting, the direct use of phage lysates for *S. aureus* treatment can compromise the purity and introduce unknown toxicity during CRISPR-Cas administration. Phage lysate toxins may arise from residual antibiotics or inducers used during phage preparation as well as naturally secreted cell products. In addition to compromising the purity of treatments, these toxins may also contribute to underestimation of PLP titers, especially in assays with a limited number of *S. aureus* cells. Therefore, we hypothesized that removal of these toxins would lead to cleaner CRISPR-Cas antimicrobials with higher apparent titers, enabling greater accuracy and control of treatments. To investigate this hypothesis, we quantified the titers of CRISPR-Cas antimicrobials directly from phage lysates and also after an additional step of ultracentrifugation.

The titering assay was performed with the standard 100 µL of filtered lysate and reduced volumes of 10, 1, and 0.1 µL, thereby diluting out the toxicity and increasing the number of *S. aureus* cells relative to PLPs. We found that phagemid titer increased as the amount of lysate used in the transduction reaction decreased (**Fig. 2F**). We also tested different MMC inducer concentrations of 2 and 0.5 µg/mL, but this variable did not have a significant impact on phagemid titer (**Fig. 2F**).

To completely remove chemical contaminants from phage preparations and further increase titer, lysates were ultracentrifuged, and phage pellets were concentrated 40x in sterile SM buffer (**Fig. 2F**). After this step, phagemid titers in excess of 10^10^ TRU/mL were consistently obtained for ∼10-kb CRISPR phagemids using only P*_terS_*-*terS* as a packaging signal. Importantly, plaque-forming units were still undetectable in purified and concentrated PLP stocks.

#### Optimized PLP production system yields high-purity CRISPR-Cas antimicrobials

We next sought to evaluate the efficacy of pure CRISPR-Cas PLPs against standard CRISPR-Cas PLPs for killing *S. aureus*. Standard PLPs were produced from the parent production strain, SaDD0001, using the *terS* gene with no promoter as a packaging signal and pure PLPs were produced from the *ΔterS,* AcrIIA4-harboring production strain, SaDD0007, using the 86-bp P*_terS_*-*terS* packaging signal (**Fig. 2E**). Despite the absence of detectable PFU from pure PLPs, phage terminases are known to erroneously package DNA[20]. Therefore, to determine the extent of unwanted DNA contamination in pure PLPs, encapsidated DNA was extracted and sequenced for both standard and pure PLP formulations and compositions were compared. In standard PLPs, we found that 92.5% of DNA was *ϕ*NM1, 7.2% was phagemid, and 0.3% was gDNA from *S. aureus* (**Fig. 2G**), which was comparable to experimentally observed TRU:PFU ratios of 8.9% for this phagemid backbone (**Fig. S2**). By contrast, pure PLPs contained 94.3% phagemid DNA, 3.2% *S. aureus* gDNA, and 2.6% *ϕ*NM1 DNA (**Fig. 2G**). Residual *ϕ*NM1 DNA in pure PLPs was highly fragmented, making it unlikely to establish phage infection. Therefore, while full-length phage genomes were excluded from pure PLPs, some contaminating DNAs were still packaged, presumably due to the nonspecificity intrinsic to the phage terminase.

Taken together, we engineered *S. aureus* SaDD0007 capable of selectively packaging CRISPR-Cas antimicrobials without self-targeting, enabling CRISPR-mediated attack on any genomic locus. Using modular phagemid cargoes harboring the minimal packaging signal and CRISPR-Cas payloads, we demonstrated SaDD0007 could produce CRISPR-Cas antimicrobials with a 94.3% purity, significantly higher than a 3% purity previously reported[8]. Ultracentrifugation can improve both the titer and purity while eliminating the toxic effects of phage lysates.

### A high-copy phagemid origin of replication drives pure PLP titers to nearly 10^12^ TRU/mL

We next sought to determine whether increasing the phagemid copy number increased the titer of CRISPR-Cas antimicrobials. To do this, the *S. aureus ori* module from the CRISPR phagemid used for pure PLPs was replaced with the *ori* from pCN56 (**Fig. 3A**), and qPCR was used to verify increased plasmid copy number (PCN). The standard, low-copy *ori* was found to have a mean PCN of 5 copies/cell, while the high-copy *ori* was maintained at 54 copies/cell, over a tenfold increase (**Fig. 3B**). PLPs were generated from strains harboring null phagemids with either low- or high-copy origins, and transducing titers were measured. High-copy PLPs achieved a mean titer of 3.20 ± 1.20×10^11^ TRU/mL, which was ∼15x higher than low-copy PLPs (**Fig. 3B**). Transductants obtained from high-copy PLPs had a small colony phenotype, suggesting increased metabolic burden (**Fig. 3C**). To our knowledge, these are the highest reported titers of pure PLPs to date and were achieved using a minimal packaging signal of 525 bp (P*_terS_*-*terS*).

**Figure 3:**
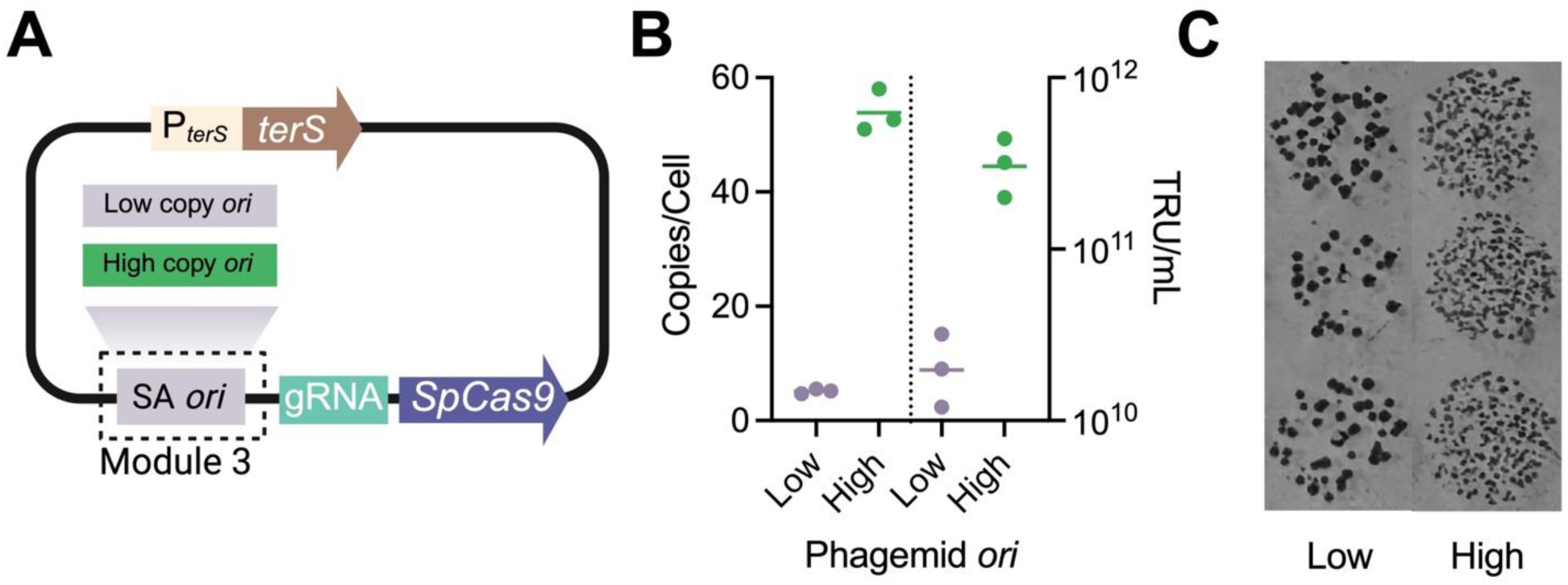
A high-copy phagemid *ori* results in elevated PLP titer and metabolic burden. (**A**) Modular phagemid design for *S. aureus* origin of replication. (**B**) Plasmid copy number and PLP titer for phagemids with low- or high-copy origins. (**C**) Colony size of transductants obtained from low- or high-copy PLPs.

### Pure CRISPR-Cas antimicrobials rely on the phage life cycle to amplify therapeutic payloads in the absence of selection

#### Activities of CRISPR-Cas antimicrobials depend on lysogeny in S. aureus

We designed a CRISPR-Cas antimicrobial with a multitargeting gRNA that can simultaneously disrupt five distinct genomic loci, increasing chromosomal damage and causing cell death (**Fig. 4A**). We performed both solid and liquid media killing assays to compare the bactericidal activities of standard and pure PLPs harboring CRISPR-Cas phagemids. First, *S. aureus* SaDD0001 was transduced with null and active PLPs, and transductants were enumerated under selection (**Fig. 4B**). Standard and pure PLPs yielded the same number of transductants, implying DNA transfer is unaffected in pure PLPs. Lethality was also conserved between formulations, with similar log_10_ reductions of 3.96 ± 0.41 and 4.11 ± 0.70 for standard and pure PLPs, respectively (**Fig. 4C**).

**Figure 4:**
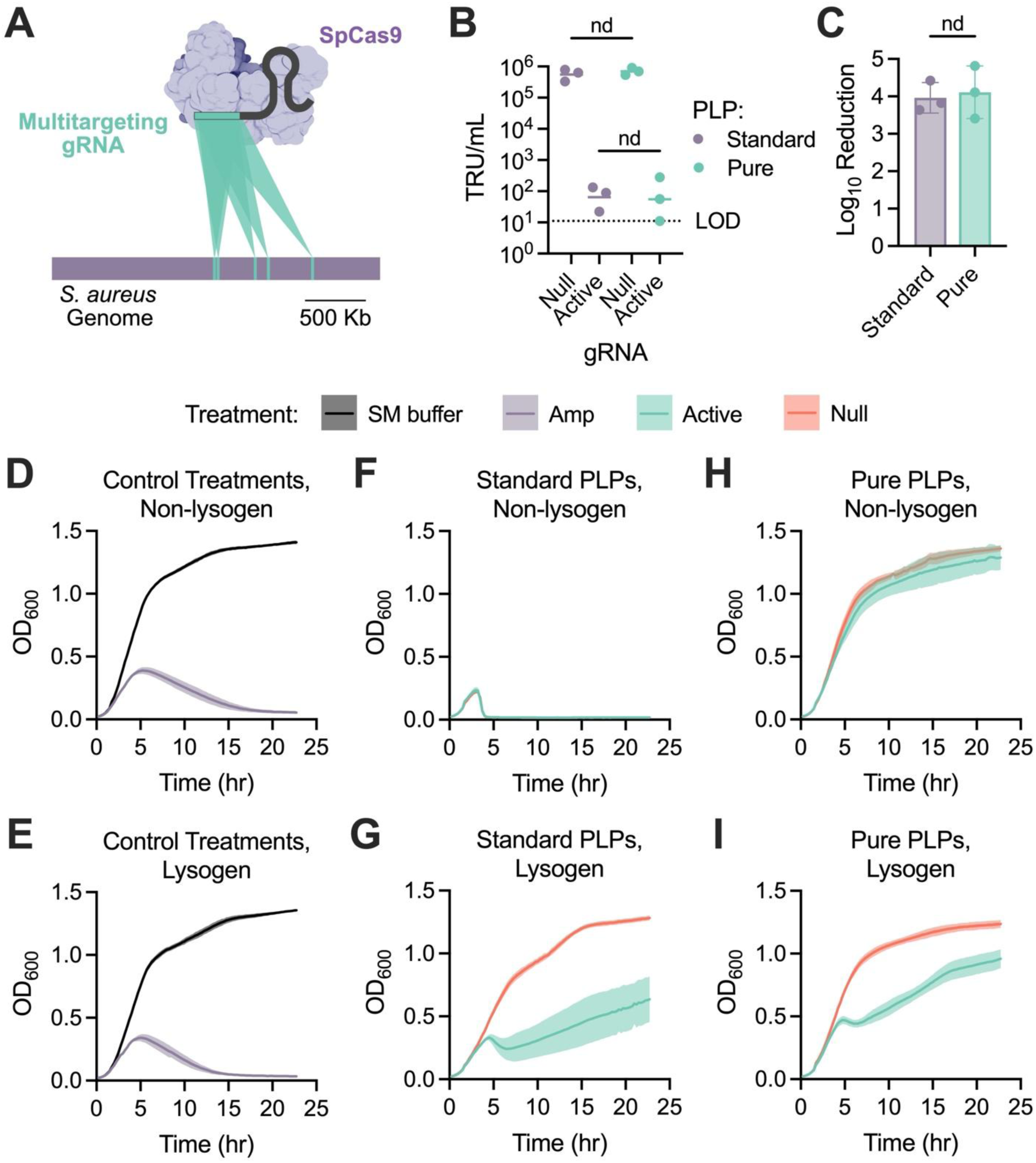
Potency of standard and pure PLPs harboring CRISPR-Cas antimicrobials on the treatment of the non-lysogen RN4420 and SaDD0001. (**A**) Visualization of the active gRNA’s target sites throughout the *S. aureus* genome. (**B**) Transduction efficiencies of standard and pure PLPs harboring either null or active CRISPR phagemids. (**C**) Log_10_ reduction of standard and pure PLPs. (**D–I**) Growth profiles of non-lysogenic or lysogenic *S. aureus* treated with (**D–E**) control treatments, (**F–G**) standard PLPs, or (**H–I**) pure PLPs. Statistical significance determined by multiple unpaired t tests assuming either (**B**) lognormal or (**C**) Gaussian distributions, where “nd” represents “not a discovery.”

Next, *S. aureus* RN4220 and SaDD0001 were infected with null and active CRISPR-Cas PLPs to evaluate killing kinetics in the absence of antibiotic selection. As controls, cells were also treated with SM buffer (negative control) and ampicillin (positive control) at its minimum bactericidal concentration[21] (MBC) (**Figs. 4D–4E**). The controls worked as expected, where ampicillin inhibited cell growth while the SM buffer did not. Standard PLPs with both null and active guides were able to kill *S. aureus* RN4220 due to its susceptibility to *ϕ*NM1 infection and the presence of contaminating *ϕ*NM1 genomes in these lysate preparations (**Figs. 2G, 4F**). In contrast, SaDD0001, whose *ϕ*NM1 prophage excludes superinfection, was inactivated in a CRISPR-dependent manner, with null PLPs demonstrating minimal toxicity and active PLPs reducing bacterial viability before eventual escape (**Fig. 4G**).

Surprisingly, pure PLPs were ineffective at killing RN4220, regardless of gRNA (**Fig. 4H**). Pure PLPs with an active guide had slightly decreased growth rate and carrying capacity relative to null treatments but failed to achieve negative growth rates at any point (**Fig. 4I**). When used to infect SaDD0001, pure PLPs displayed similar killing signatures as standard PLPs, but with reduced efficacy (**Fig. 4I**). Since the only genetic difference between RN4220 and SaDD0001 is the presence of the *ϕ*NM1 prophage, these results led us to conclude that certain *ϕ*NM1 proteins are necessary for the CRISPR antimicrobial activity of pure PLPs in the absence of selection.

#### The prophage of lysogenic S. aureus induces self-replication of pure CRISPR-Cas antimicrobials

We hypothesized that without selection, therapeutic phagemids delivered by pure PLPs require participation in the phage life cycle to persist in a cell population. To test this hypothesis, we repeated the liquid killing assay for pure PLPs carrying either null or active guides and sampled at regular intervals to enumerate phagemid and phage genome titers as indicated by TRU/mL and PFU/mL, respectively (**Fig. 5A**). Null PLPs behaved similarly across both strains, while active PLPs had strain-specific activity (**Fig. 5B**). As before, CRISPR lethality was more pronounced in SaDD0001 than in RN4220, achieving higher growth rate reduction and lower final OD_600_ (**Fig. 5B**). Consistent with our hypothesis, phagemid titer gradually decreased over time for both null and active PLPs targeting RN4220, despite all samples starting at the same concentration (**Fig. 5C**). Conversely, SaDD0001 treated with active, but not null, PLPs showed an increase in phagemid concentration over the first three hours of infection before reaching a plateau at ∼5×10^7^ TRU/mL (**Fig. 5C**). Samples obtained from RN4220 had almost no detectable plaque-forming units across infection (**Fig. 5D**). Surprisingly, however, at five and six hours post-infection, extremely low phage titers were observed for both active and null PLPs (**Fig. 5D**). This indicates that whole phage genomes are still being packaged into pure PLPs at nearly undetectable levels yet remain capable of amplifying during infection. Phage titers recovered from SaDD0001 treated with null PLPs remained constant across infection at ∼5×10^4^ PFU/mL, consistent with spontaneous prophage induction observed for other staphylococcal phages[22] (**Fig. 5D**). For SaDD0001 treated with active PLPs, phage titer quickly increased in the first three hours after infection, then remained steady at ∼5×10^8^ TRU/mL, mirroring the trend observed for phagemid titers (**Fig. 5D**). Across all time points, phagemid and phage titers were tightly correlated, with an R^2^ = 0.81 (**Fig. S3**).

**Figure 5:**
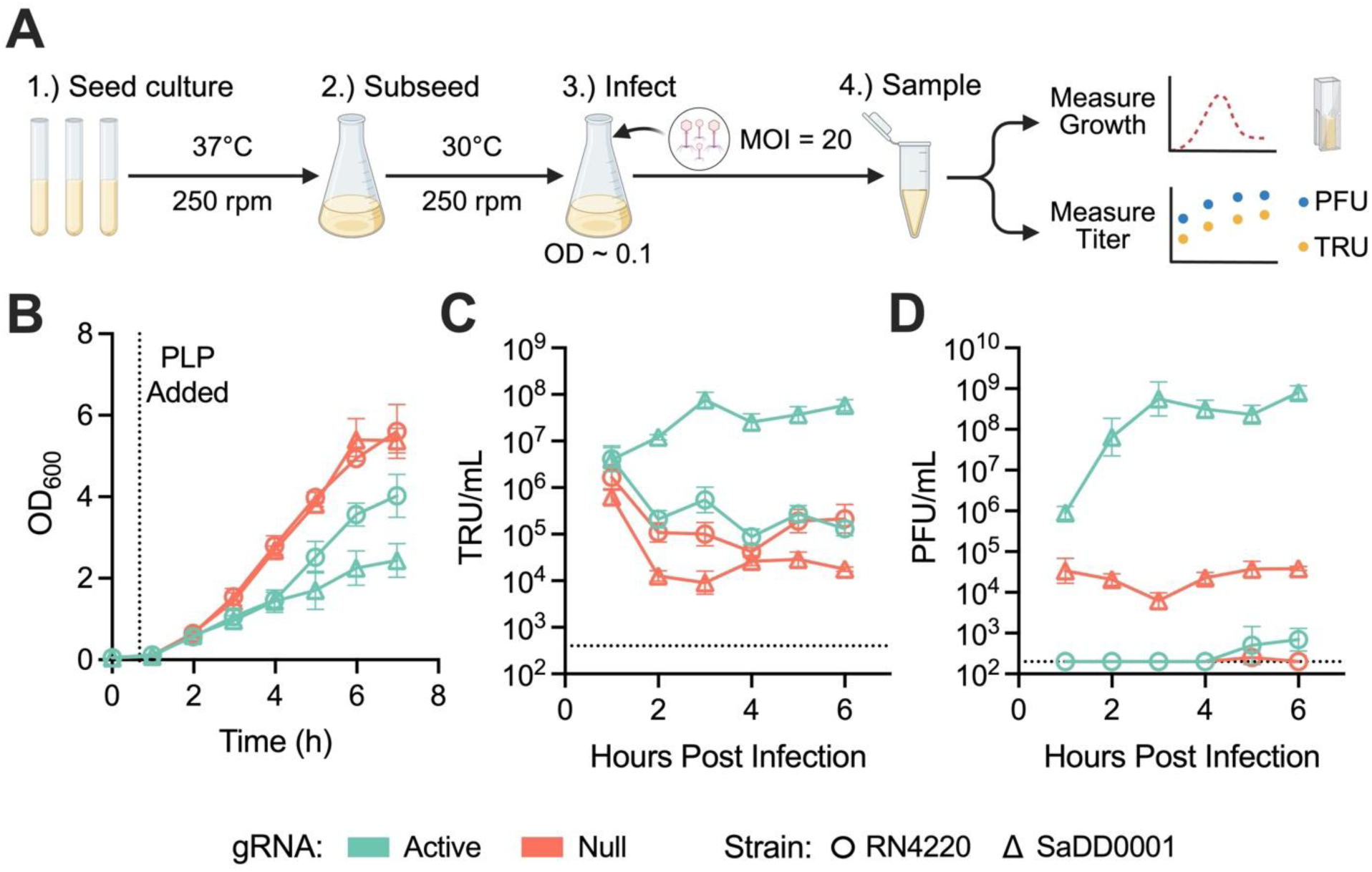
CRISPR PLPs rely on the phage life cycle to amplify therapeutic titer in the absence of selection. (**A**) Workflow for titer tracking assay. (**B**) Growth curves of *S. aureus* cultures treated with pure PLPs harboring either an active or null CRISPR system. (**C–D**) Titers of (**C**) therapeutic PLPs or (**D**) phages following infection of lysogenic *S. aureus* with either active or null pure PLPs.

These data show that CRISPR antimicrobial activity is impacted by the concentration of therapeutic PLPs in the cell population, and that, in turn, the titer of transducing PLPs is affected by participation in the phage life cycle (**Figs. 5B, 5C**). CRISPR-induced DNA damage has previously been shown to induce the SOS response and resident prophages[23, 24]. As the prophage excises and replicates, both WT phage and therapeutic PLPs are produced, which are then able to infect new cells after host lysis. This model explains why only an active gRNA was able to trigger PLP and phage formation, since an NT gRNA does not induce DNA damage (**Figs. 5C–5E**). These findings are relevant for informing the design of next-generation CRISPR antimicrobials, as phage DNA in CRISPR PLPs has typically been viewed as an undesirable contaminant. Nevertheless, pure PLPs may still be useful for treating lysogenic pathogens—for example, *S. aureus*, which is known to commonly possess one or more prophages[25, 26].

## Conclusion

In this study, we define a simple and streamlined system for generating high-purity and high-titer phage-like particles. We reveal that a packaging signal of *terS* under its native promoter is sufficient and minimal for efficient phagemid encapsidation and exclusion of wild-type phage genomes in a Δ*terS* production strain. Packaging efficiency was greatly reduced when *terS* expression was driven by a strong constitutive promoter, indicating that precise expression of the terminase protein is crucial for favorable packaging outcomes, in addition to the presence of the *pac* site. Despite vast improvements in purity, low levels of contaminating phage DNA could be detected in pure PLPs via sequencing and following amplification in a prophage-deficient strain. Since the cognate *pac* site was deleted from the production strain, this spurious packaging likely arises from the nonspecificity inherent to phage terminases, as seen in generalized transduction. This limitation could potentially be addressed by protein engineering of the terminase for improved affinity to its cognate *pac* site or through removal of pseudo-*pac* sites throughout the production strain.

CRISPR antimicrobials generated using this approach rely on phage biology to amplify therapeutic payloads in the absence of selection. This finding challenges the traditional stance that phage DNA is an unwanted contaminant in CRISPR antimicrobials, instead suggesting it may act as an adjuvant by spreading therapeutic phagemids across a treated population. As such, efforts toward detoxifying co-resident phages, instead of completely removing them, could be warranted. Admittedly, a limitation of this study is that we did not determine whether increased dosage could offset the inability of pure PLPs to replicate; however, using a high-copy *ori*, we raised PLP titers to levels that enable this hypothesis to be directly tested in future work. Collectively, these results simplify the genetic requirements for efficient packaging in *pac*-type phages, clarify the role of phage DNA in CRISPR PLP activity, and provide valuable insights for designing new phage-based treatments with improved purity, tunability, and translational potential.

## Materials and Methods

### Culture Conditions and Strains

The plasmids, strains, and primers used for this study can be found in **Supplementary Tables S1**, **S2**, and **S3**, respectively. The base plasmid used for packaging gene complementation vectors was pRMC2 (Addgene #68940) and was a gift from Tim Foster[27]. The base plasmid used for CRISPR antimicrobial phagemids was pCasSA (Addgene #98211) and was a gift from Quanjiang Ji[3]. AcrIIA4 was sourced from pCSW1 (Addgene #86836)[28] and integrated into *S. aureus* using a derivative of pRN111 (Addgene #84463)[29]. These plasmids were gifts from Joseph Bondy-Denomy and Reindert Nijland, respectively.

Unless mentioned otherwise, all strains were cultured at 37 °C, 250 rpm. Lysogeny broth (LB) with 50 µg/mL kanamycin or 100 µg/mL ampicillin, as appropriate, was used to culture *E. coli* strains. Tryptic soy broth (TSB) was used to culture *S. aureus* and 10 µg/mL chloramphenicol was used for selection, when required. For phage infection experiments, CaCl_2_ was added to a final concentration of 5 mM and dipotassium phosphate was replaced with 5 g/L MOPS to prevent calcium phosphate precipitation. When solid agar was required, 1.5% LB agar was used for *E. coli,* and 1.5% tryptic soy agar (TSA) was used for *S. aureus*. Soft agar for plaque assays was prepared as 0.7% TSA.

### Molecular Cloning

Plasmids were either constructed with *E. coli* 10-beta (NEB #C3019H), Clean Genome LowMut (Scarab Genomics #C-6786), or CopyCutter™ EPI400™ (LGC Biosearch Technologies #C400EL10). PCR fragments were amplified using Phusion™ Hot Start II DNA Polymerase (Thermo Scientific™ #F549L) according to the manufacturer’s recommendations. All cloning reactions were performed at a final volume of 10 µL. Ligation reactions employed 0.5 µL NEB T4 DNA ligase (#M0202) and were incubated overnight at 16 °C. Gibson assemblies were performed as previously described^[30]^. Golden Gate assemblies utilized 0.5 µL NEB BbsI-HF (#R3539) or BsaI-HFv2 (#R3733) and 0.5 µL T4 ligase. Golden Gate reactions were cycled 60 times between 37 °C and 16 °C for 5 minutes each, then incubated 1 hour at 37 °C, 20 minutes at 80 °C, and held at 4 °C until used. When annealed oligos were used for cloning, oligos were first phosphorylated by combining 1 µL of forward and reverse oligo with 1 µL NEB T4 polynucleotide kinase (#M0201), 5 µL NEB T4 ligase 10x buffer (#B0202), and 42 µL Millipore water and incubating for an hour at 37 ℃. After phosphorylation, 2.5 µL of 1M NaCl was added and oligos were annealed at 95 ℃ for 5 minutes and allowed to slowly cool to room temperature. Annealed oligos were stored at 4 ℃ until used. Plasmids were transformed into *E. coli* using heat shock and into *S. aureus* using electroporation, as previously described[31, 32]. All constructs were confirmed by colony PCR and Sanger or whole plasmid sequencing.

#### Packaging gene complementation vectors

A base shuttle vector, pDD1004, was fashioned using pRMC2 as a backbone. To assemble pDD1004, three amplicons were combined in a Gibson assembly reaction: DD001.f/r amplified from pRMC2, DD002.f/r amplified from pCasSA, and DD003.f/r amplified from pCSW21. The empty phage complementation vector, pDD0022, was prepared by digesting pDD1004 with XbaI and SalI and ligating in annealed oligos DD004.f/r, which contained a BbsI stuffer. All complementation vectors were cloned using Golden Gate assembly with BbsI-HF except for pDD1038 (*terS*), which was cloned using Gibson assembly. Packaging genes were amplified from *S. aureus* SaDD0001 gDNA using the following primer pairs: i.) DD005.f/r for pDD1038 (*terS*), ii.) DD006/DD007 for pDD1040 (P*_rpsL_*-*terS*), iii.) DD008/DD009 for pDD1043 (P*_terS_*-*terS*), iv.) DD0010/DD0011 for pDD2005 (*terS*-*terL*), v.) DD006/DD0011 for pDD2006 (P*_rpsL_*-*terS*-*terL*), and vi.) DD008/DD0012 for pDD2007 (P*_terS_*-*terS*-*terL*). Plasmid backbones were either amplified or used directly in Golden Gate assembly. For plasmids pDD1038, pDD1040, pDD2005, and pDD2006, the pDD1004 backbone was amplified using primer pairs DD0013.f/r, DD0014/DD0015, DD0016/DD0017, and DD0016/DD0015, respectively. For plasmids, pDD1043 and pDD2007, pDD0022 was used directly as the backbone. The full 186-bp *terS* promoter and *terS* gene were amplified from SaDD0001 gDNA using DD0018/DD009 and inserted directly into pDD0022 using Golden Gate assembly to create pDD1055.

#### TerS knockout vector

To delete *terS*, the pCasSA editing vector was used[3]. To begin, ∼500-bp homology arms upstream and downstream of *terS* were amplified from *S. aureus* SaDD0001 gDNA using DD0019.f/r and DD0020.f/r, respectively. Homology arm amplicons were concatenated using overlap extension PCR. Briefly, 1 µL of unpurified amplicons were combined with 37.2 µL Phusion master mix and self-amplified for 15 cycles using an annealing temperature equal to the overlapping region of the fragments. After 15 cycles, 0.4 µL of each outside primer (DD0019.f and DD0020.r) was added and the reaction was amplified for another 30 cycles with the annealing temperature set to the lowest melting temperature of the primer pair. The combined homology arms were purified and ligated into pCasSA digested with XbaI and XhoI to create pDD1030. A 20-bp guide RNA targeting *terS* was designed with CASPER[33] and ordered as complementary forward and reverse oligos possessing BsaI-compatible overhangs when annealed (SA_terS.f/r). Oligos were phosphorylated, annealed, and inserted into pDD1030 using BsaI-mediated Golden Gate assembly as previously described[31]. The final plasmid, pDD1030-1, was used to generate the *terS* deletion strain.

#### CRISPR antimicrobial phagemids

An in-house phagemid, pDD0021[31], was used as the starting point for subsequent CRISPR antimicrobial phagemids in this study. This plasmid was constructed from the pCasSA backbone and includes a non-temperature-sensitive *S. aureus ori* from pSGFPS1 and a *terS* packaging signal. To enable BsaI-mediated Golden Gate assembly, a BsaI recognition site was removed from the *S. aureus ori* by introducing mutations with primers. Two fragments from pDD0021 were amplified with DD0021.f/r and DD0022.f/r and combined using Gibson assembly. The resultant phagemid, pDD0023, served as the backbone for all standard PLP preparations. To create the base phagemid for pure PLPs, pDD0029, *terS* was replaced with the P*_terS_*-*terS* packaging signal from pDD1043. P*_terS_*-*terS* was amplified from pDD1043 with DD0023.f/r and inserted into pDD0023 digested with XbaI and XhoI using Gibson assembly.

To create active phagemids, a self-targeting gRNA was inserted into either pDD0023 or pDD0029 using Golden Gate assembly with annealed oligos SA_active.f/r, as previously described[31]. Null phagemids were similarly created using a non-targeting gRNA from annealed oligos SA_null.f/r.

#### High-copy phagemid variants

To create a high-copy phagemid variant, the backbone of pDD0021 was amplified with DD0024.f/r and the *S. aureus ori* from pCN56 was amplified with DD0025.f/r. These fragments were combined using Gibson assembly to create pDD0027. Once the plasmid copy number was verified for pDD0027, another high-copy phagemid, pDD0034, was constructed with the same primers as before, but with pDD0029 as the backbone instead of pDD0021. This allowed for direct comparison of packaging efficiency in pure PLPs.

### AcrIIA4 Integration into *S. aureus*

To create the CRISPR antimicrobial production strains, AcrIIA4 was integrated into either *S. aureus* SaDD0001 or SaDD0003 using pDD1044, an in-house vector based on pRN111[29, 31]. In short, pDD1044 was transformed into *S. aureus* and transformants were restreaked on selective agar and grown overnight at 45 ℃. This process was repeated once more. Colonies were then inoculated into non-selective media and cultured at 30 ℃ for 2.5 days. Each morning and each evening, cultures were back diluted 1000x into fresh media. After 2.5 days, cultures were serially diluted in sterile Millipore water, plated on TSA containing 100 ng/mL anhydrotetracycline, and incubated overnight at 37 ℃. Insertion mutants were screened for using colony PCR and verified for plasmid curing by restreaking on selective and non-selective agar. Mutants lacking the editing plasmid were cultured and their gDNA was extracted, as previously described[32]. AcrIIA4 insertion was confirmed by PCR and Sanger sequencing.

### *TerS* Knockout

To generate the *ΔterS S. aureus* strain, 1 µg of pDD1030-1 was transformed into *S. aureus* SaDD0001, as previously described[31]. Transformants were screened with colony PCR using primers DD0026.f/r to identify knockout mutants and confirmed colonies were inoculated into 1 mL TSB and grown overnight at 30 ℃. Overnight cultures were diluted 1000x into fresh media and incubated at 42 ℃ for ∼9 hours to cure the editing plasmid. Afterwards, cultures were restreaked on TSA and incubated overnight at 37 ℃. Colonies were again screened via colony PCR to confirm the *terS* knockout and also restreaked on both selective and nonselective TSA to verify the loss of plasmid. Colonies with both a *terS* knockout and cured plasmid were inoculated into 3 mL TSB and grown overnight at 37 ℃. Genomic DNA was extracted and amplified with DD0027.f/r, which flank the *terS* deletion site. Resultant amplicons were Sanger sequenced with DD0027.f to confirm the knockout phenotype.

### Phages and Phage-Like Particles

#### Phage and PLP generation

All phages and phage-like particles were generated using chemical induction. When phage-like particles were generated, production strains were cultured with antibiotic selection to ensure phagemid retention. Production strains were seeded into 1 mL TSB and grown overnight. Overnight cultures were diluted 50x into fresh media and grown to mid-exponential phase. Mid-exponential cultures were induced with either 2 or 0.5 µg/mL MMC and transferred to 30 ℃ for 16–20 hours. Lysed cultures were centrifuged at 3,000 xg for 10 minutes and the supernatant was passed through a sterile 0.22 µm filter. When phages or PLPs were further concentrated, filtered lysates were spun down for 40 minutes at 35,000 xg and 4 ℃. Supernatants were discarded and phage pellets were resuspended in 1/40^th^ of the initial volume using SM buffer.

#### Phagemid titer quantification

When CRISPR elements were present on the phagemids to be titered, the recipient strain was *S. aureus* SaDD0006[31], which contains AcrIIA4 and is resistant to SpCas9 activity. Otherwise, *S. aureus* SaDD0001 was used. The recipient strain was grown overnight in 1 mL TSB supplemented with 5 mM CaCl_2_. Overnight cultures were diluted 50x into fresh media and grown until OD_600_ ≥ 1. The recipient culture was diluted back to exactly OD_600_ = 1.0 and 100 µL was added to a sterile 96-well microplate. For phage complementation experiments, 100 µL of filtered phage lysate was mixed with the culture. When lower volumes of phage lysate were used (e.g., for highly concentrated lysates), media was added to the lysate to achieve a final volume of 100 µL before addition to the microplate. Transduction reactions were allowed to incubate statically for 1 hour at 37 ℃. After incubation, transduction reactions were serially diluted and spotted on selective agar either by hand or with an Opentrons Flex liquid handling robot. Spots were allowed to dry for 5–10 minutes in the biological hood before incubation overnight. The next day, colonies were enumerated and used to back calculate the transducing titer of the phage lysate according to the equation below.

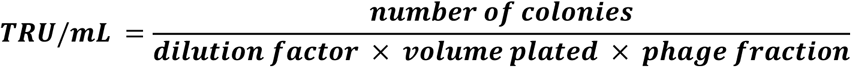

#### Plaque assays

*S. aureus* RN4220 was seeded into 1 mL TSB with 5 mM CaCl_2_ and grown overnight. The overnight culture was diluted 50x into fresh media and allowed to grow to mid-exponential phase. In a sterile culture tube, 4.5 mL of 0.7% TSA with 5 mM CaCl_2_ was mixed with 500 µL of culture and poured over a 1.5% TSA petri dish. The soft agar was allowed to solidify in the biological hood for ∼5 minutes before serial dilutions of the phage lysate were spotted. Spots were allowed to dry and then plates were incubated at 37 ℃ overnight with the agar side up. The following morning, plaques were enumerated and used to calculate the plaque-forming titer of the phage lysate.

#### PLP DNA extraction and sequencing

To aid in stabilization and DNA precipitation, 12.5 μL of 1 M MgCl_2_ was added to 1 mL of high titer (>10^9^ PFU/mL) PLP lysate. The lysate was gently mixed, then 4 μL of DNAse I (25 mg/mL) and 1 μL of RNase A (100 mg/mL) were added. After briefly vortexing, the mixture was incubated for 30 minutes at room temperature. In this order, 50 μL of 0.5 M ethylenediaminetetraacetic acid (EDTA), 0.63 μL of Proteinase K (20 mg/mL), and 50 μL of 10% SDS were added. The mixture was vortexed vigorously, then incubated for 60 minutes at 55 ℃. During incubation, the mixture was vortexed at 20-minute intervals. The final mixture was split into two 500 μL aliquots, and 500 μL of 25:24:1 phenol-chloroform-isoamyl alcohol (PCI) was added to purify the mixture of residual proteins and contaminating materials. The sample was inverted several times, then centrifuged for 5 minutes at 13,000 rpm for removal of the top aqueous layer. This was repeated until no white interphase remained, taking care to avoid phenol contamination. To precipitate the DNA, 1 mL of 95% ethanol and 50 μL of 3 M sodium acetate were added to the aqueous layer, then incubated overnight at −80 ℃. The sample was gently mixed until a pellet formed, then centrifuged for 10 minutes at 13,000 rpm. The pellet was washed with 500 μL of 80% ethanol, then air dried at 50 ℃ for 10 minutes. Finally, the pellet was dissolved in 50 μL of Millipore water, then incubated at 37 ℃ for 30 minutes to ensure complete solvation. Purified DNA was sequenced with Nanopore sequencing by Eurofins Genomics.

#### Solid media killing assays

Solid media killing assays were performed as previously described[31]. In short, the target strain, *S. aureus* SaDD0001, was grown overnight in 1 mL TSB with 5 mM CaCl_2_. The overnight culture was diluted 50x into fresh media, then incubated until OD_600_ 0.4–0.6. The culture was back diluted to exactly OD_600_ = 0.04 and 100 µL was aliquoted into a sterile 96-well microplate. PLPs were added at an MOI = 1 based on phagemid titer in a volume of 100 µL, and the reaction was allowed to incubate statically at 37 ℃ for 1 hour. After 1 hour, 180 µL of the transduction reaction was plated on selective TSA and incubated overnight. Surviving colonies were enumerated and the TRU/mL for active and null treatments were calculated. Log_10_ reduction was calculated using the equation below.

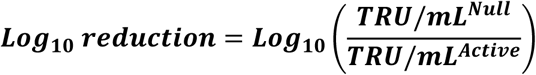

#### Microplate killing assays

Microplate killing assays were performed as previously described[31]. *S. aureus* RN4220 and SaDD0001 were grown overnight in TSB with 5 mM CaCl_2_. Cultures were diluted 50x in fresh media and grown to OD ∼ 0.4–0.5 at 30 ℃. Cultures were normalized to exactly OD_600_ = 0.05 and 190 µL was aliquoted into a sterile 96-well microplate. The plate was incubated at 30 ℃ with double orbital shaking, a temperature gradient of 1 ℃ to prevent condensation, and 15 minute read intervals, in an Agilent BioTek Epoch 2 plate reader until OD_600_ ∼ 0.1. 10 µL of treatments or SM buffer were added, and the plate was returned to the plate reader for ∼23 hours of incubation. PLP lysates were added at an MOI = 20 and ampicillin was added at a final concentration equal to its MBC, 6 µg/mL[21].

#### Titer tracking assays

Target strains were seeded and subcultured in the same manner as for microplate killing assays. Exponential cultures were diluted to OD_600_ = 0.05 and 10 mL was aliquoted into 50 mL baffled Erlenmeyer flasks. Flasks were incubated at 30 ℃ until OD_600_ = 0.1, at which point PLPs were added at an MOI = 20. Every hour for the next 6 hours, 500 µL samples were taken and immediately processed as follows. OD_600_ was measured using 100 µL of sample, and the remaining sample was centrifuged at 4,700 xg for 5 minutes. Supernatants were transferred to a sterile 96-well plate, which was sealed and kept at 4 ℃ until the next round of samples. After all samples were collected, supernatants were filtered through a 0.22 µm membrane and lysates were titered for TRU/mL and PFU/mL, as described above.

### Plasmid Copy Number Determination

Quantitative PCR (qPCR) was used to calculate the plasmid copy number (PCN) of pDD0027 (pCN56 *ori*) compared to pDD0021 (pSGFPS1 *ori*). *S. aureus* SaDD0001 harboring either pDD0021 or pDD0027 was grown in selective TSB overnight and diluted 50x into fresh media on the following day. Overnight cultures were diluted to OD_600_ = 0.05 in 2 mL fresh media and grown to mid-exponential phase. Mid-exponential cells were spun down at 6,000 xg for 3 minutes and resuspended in 100 µL sterile Millipore water containing 50 µg/mL lysostaphin. Cells were incubated in a water bath at 37 ℃ for 15 minutes to allow lysis to occur. Total DNA was then extracted from lysed cells using a ZymoBIOMICS DNA/RNA Miniprep Kit (#R2002), per manufacturer’s recommendations. DNA concentration was measured using a Nanodrop and qPCR reactions were set up to determine the PCN.

To perform qPCR, the Qiagen QuantiTect SYBR Green PCR kit (#204143) was used with an Applied Biosystems™ StepOnePlus™ Real-Time PCR System. Reactions were performed in triplicate for 5 template DNA concentrations and 2 qPCR probes. DNA concentrations ranged from 10 ng/µL to 1 pg/µL in serial dilutions, while probes amplified ∼180-bp regions from i.) the single-copy *nuc* gene in the *S. aureus* chromosome (DD0028.f/r) or ii.) SpCas9 found in both pDD0021 and pDD0027, but not *S. aureus* (DD0029.f/r). No template controls were included for each probe. Each qPCR reaction contained 10 µL 2x SYBR Green master mix, 8 µL DNA, 1.8 µL Millipore water, and 0.1 µL each primer at 100 µM concentration. Reactions were cycled according to Qiagen’s recommendations using a melting temperature of 58 ℃. Using the resultant Ct values and known DNA concentrations, PCN was calculated as previously described[34].

### Computational Analyses

#### Encapsidated DNA composition

Adapters were removed from raw Nanopore reads using porechop_abi with the -abi flag[35]. Reads were then filtered using filtlong (--min_length 200 --keep_percent 90) to retain the highest quality reads[36]. Preprocessed reads were mapped to reference sequences using minimap2 with the long-read high-accuracy preset (-x lr:hq), which optimizes alignment performance for long-read sequencing data[37]. SAM-formatted alignments were converted to sorted BAM files using samtools (view -bS and sort commands), and sorted BAM files were indexed for efficient downstream analysis. Alignments were filtered to exclude secondary and supplementary alignments using samtools (view -F 2304) and aligned reads were tallied per reference. The number of aligned bases was calculated by parsing CIGAR strings and summing reference-consuming operations (M, =, X, and D). Unmapped reads were excluded from downstream analyses, and relative abundance was calculated as the proportion of total bases mapping to each reference.

#### Statistical analyses

All statistical tests were performed in Graphpad Prism v10.6.1 with default settings.

## Author contributions

Conceptualization: C.T.T and D.S.D. Methodology: D.S.D and H.B. Resources: C.T.T. Investigation: D.S.D, H.B., and C.T.T. Formal analysis: D.S.D, H.B., and C.T.T. Validation: D.S.D, H.B., and C.T.T. Visualization: D.S.D and C.T.T. Funding acquisition: C.T.T. Supervision: C.T.T. Writing—original draft: D.S.D. and C.T.T. Writing—review and editing: D.S.D. and C.T.T.

## Acknowledgements

The authors would like to sincerely thank Dr. Ed Wright and the Bioanalytical Resource Facility at the University of Tennessee for access to the Beckman Optima MAX-XP ultracentrifuge. The following reagents were obtained through BEI Resources, NIAID, NIH: *Staphylococcus aureus* Fluorescent Reporter Plasmid pSGFPS1, Recombinant in *Staphylococcus aureus*, NR-51163. The following reagent was provided by the Network on Antimicrobial Resistance in Staphylococcus aureus (NARSA) for distribution by BEI Resources, NIAID, NIH: Escherichia coli – Staphylococcus aureus Shuttle Vector pCN56, Recombinant in Escherichia coli, NR-46156.

