## Supplementary Figures S1-S3 and Tables S1-S3 for "A Minimal Packaging Signal Enables Production of High-Purity Phage-Like Particles for CRISPR-Cas Antimicrobials in *Staphylococcus aureus*"

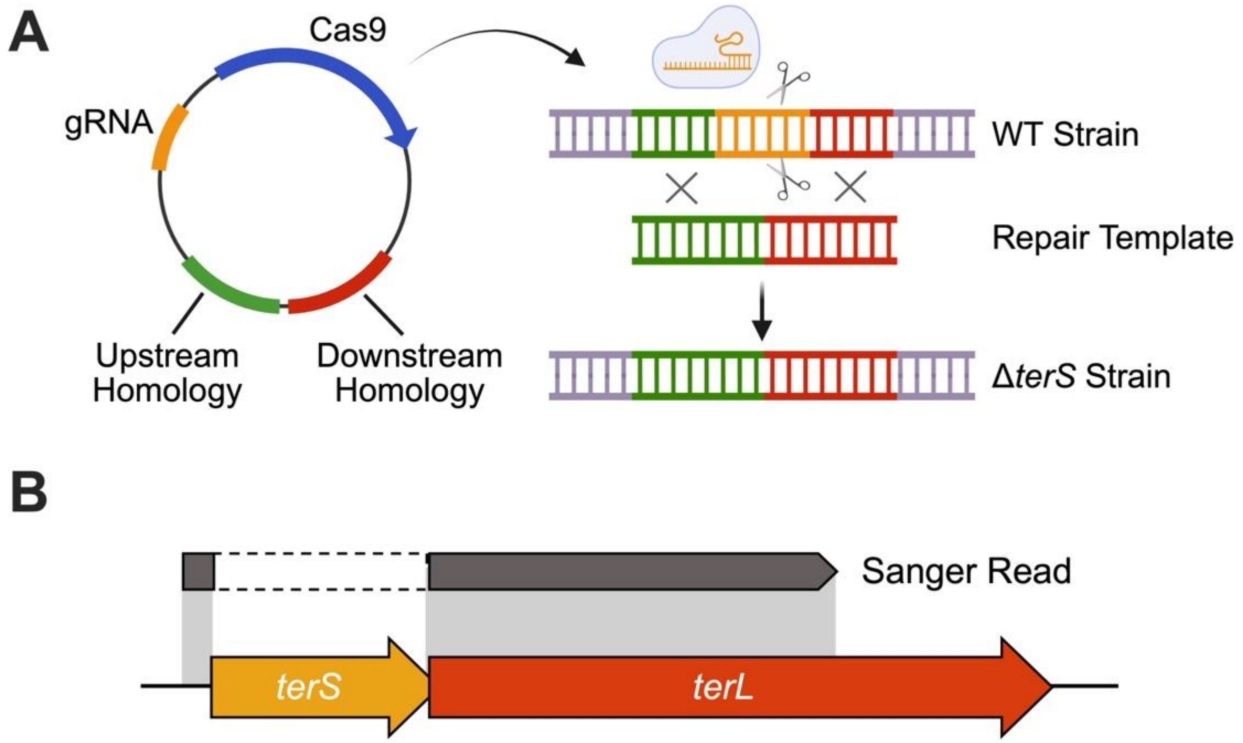


**Figure S1:** Generation of *terS* knockout. (**A**) CRISPR editing workflow for *terS* knockout. (**B**) Sanger sequencing read alignment for *terS* deletion strain.

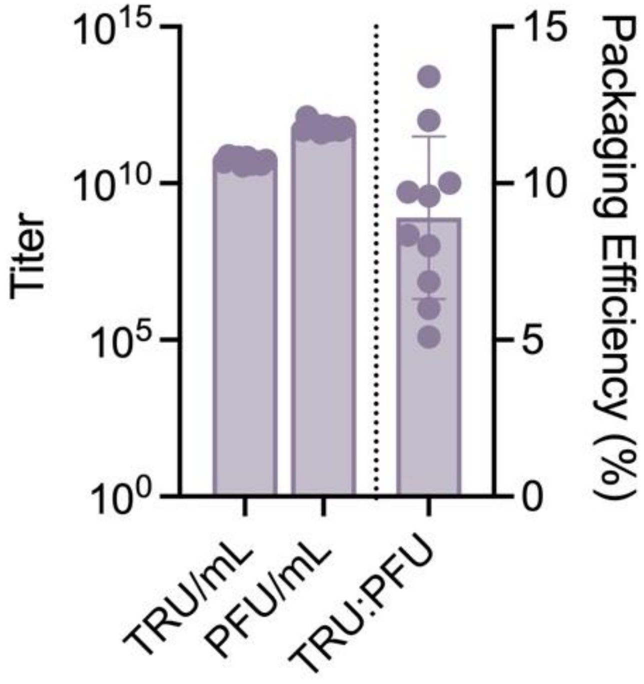


**Figure S2:** Transducing titer, plaque-forming titer, and packaging efficiency for standard PLPs harboring ~10 kb phagemids.

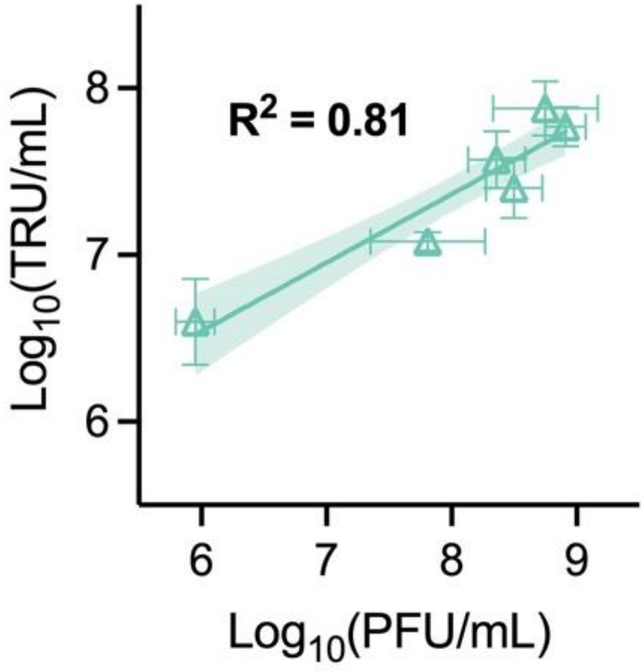


**Figure S3:** Correlation of log-transformed TRU/mL with log-transformed PFU/mL for lysogenic *S. aureus* treated with active pure PLPs. Shaded band represents the 95% confidence interval.

**Table S1:** List of plasmids used in this study.

| Plasmid ID | Parent Plasmid | Role in Study | gRNA | Source |
|---|---|---|---|---|
| pRMC2 | | Base plasmid | | Addgene #68940 |
| pCasSA | | Base plasmid | | Addgene #98211 |
| pCSW21 | | AcrIIA4 source | | Addgene #86836 |
| pRN111 | | Base plasmid | | Addgene #84463 |
| pSGFPS1 | | Non-ts *S. aureus ori* source | | BEI (NR-51163) |
| pCN56 | | High-copy E. coli and S. aureus shuttle vector | | BEI (NR-46156) |
| pDD0021 | | SpCas9 phagemid w/ non-ts *S. aureus ori* | | Dooley, Trinh 2026 |
| pDD0023 | | Base SpCas9 phagemid | | Dooley, Boyd, Trinh 2026 |
| pDD1044 | pRN111 | AcrIIA4 integrative vector | | Dooley, Trinh 2026 |
| pDD1004 | pRMC2 | Base shuttle vector | | This Study |
| pDD0022 | pDD1004 | Empty phage complementation vector | | This Study |
| pDD1038 | pDD1004 | Phage complementation vector (*terS*) | | This Study |
| pDD1040 | pDD1004 | Phage complementation vector ($P_{rpsL}$-*terS*) | | This Study |
| pDD1043 | pDD0022 | Phage complementation vector ($P_{terS}$-*terS*) | | This Study |
| pDD2005 | pDD1004 | Phage complementation vector (*terS*-*terL*) | | This Study |
| pDD2006 | pDD1004 | Phage complementation vector ($P_{rpsL}$-*terS*-*terL*) | | This Study |
| pDD2007 | pDD0022 | Phage complementation vector ($P_{terS}$-*terS*-*terL*) | | This Study |
| pDD1055 | pDD0022 | Phage complementation vector ($P_{full\ terS}$-*terS*-*terL*) | | This Study |
| pDD1030 | pCasSA | pCasSA with homology arms for *terS* knockout | | This Study |
| pDD1030-1 | pDD1030 | Final vector for *terS* knockout | SA_terS | This Study |
| pDD0029 | pDD0023 | Pure SpCas9 phagemid ($P_{terS}$-*terS*) | | This Study |

| | | | | |
|---|---|---|---|---|
| pDD0023-active | pDD0023 | Base SpCas9 phagemid with active gRNA | SA_active | This Study |
| pDD0023-null | pDD0023 | Base SpCas9 phagemid with null gRNA | SA_null | This Study |
| pDD0029-active | pDD0029 | Pure SpCas9 phagemid with active gRNA | SA_active | This Study |
| pDD0029-null | pDD0029 | Pure SpCas9 phagemid with null gRNA | SA_null | This Study |
| pDD0027 | pDD0021 | High-copy variant of pDD0021 | | This Study |
| pDD0034-active | pDD0029-active | Pure, high-copy SpCas9 phagemid with active gRNA | SA_active | This Study |
| pDD0034-null | pDD0029-null | Pure, high-copy SpCas9 phagemid with null gRNA | SA_null | This Study |

**Table S2:** List of strains used in this study.

| Strain | Parent Strain | Relevant Features | Source |
|---|---|---|---|
| *E. coli* 10-beta | *E. coli* DH10β | | NEB (C3019H) |
| *E. coli* Clean Genome LowMut | | IS element-deficient | Scarab Genomics (C-6786) |
| *E. coli* CopyCutter™ EPI400™ | | inducible *pcnB* for copy number control | LGC Biosearch Technologies (#C400EL10) |
| *S. aureus* RN4220 | | prophage-deficient, restriction-deficient, *agrA* mutant | BEI (NR-45946) |
| *S. aureus* SaDD0001 | *S. aureus* RN4220 | single $\phi$NM1 lysogen | Dooley, Trinh 2026 |
| *S. aureus* SaDD0006 | *S. aureus* SaDD0001 | single $\phi$NM1 lysogen, $P_{SarAP1}$-*AcrIIA4*-$T_{lambda}$ inserted between NWMN_0029/0030 | Dooley, Trinh 2026 |
| *S. aureus* SaDD0003 | *S. aureus* SaDD0001 | single $\phi$NM1 lysogen, Δ*terS* | This Study |
| *S. aureus* SaDD0007 | *S. aureus* SaDD0006 | single $\phi$NM1 lysogen, Δ*terS*, $P_{SarAP1}$-*AcrIIA4*-$T_{lambda}$ inserted between NWMN_0029/0030 | This Study |

**Table S3:** List of primers used in this study.

| Name | Sequence | Purpose |
|---|---|---|
| DD001.f | TAAAAAGTGAGTTGAACTAACACGGTAACCAAGATGTCGAG | Forward for amplifying pRMC2 backbone |
| DD001.r | GAGCATTCTAGACCATGGGTGCTGGCGTAATAGCGAAG | Reverse for amplifying pRMC2 backbone |
| DD002.f | ACCCATGGTCTAGAATGCTCG | Forward for amplifying pRpsL from pCasSA |
| DD002.r | TTGATTTCTCTAATTAAGTCATTAATATTCATTCCATGTGATATGTCCTCCTCTC | Reverse for amplifying pRpsL from pCasSA |
| DD003.f | ATGAATATTAATGACTTAATTAGAGAAATCAAAAAC | Forward for amplifying AcrIIA4 from pCSW21 |
| DD003.r | TTAGTTCAACTCACTTTTTAAGGTGATT | Reverse for amplifying AcrIIA4 from pCSW21 |
| DD004.f | CTAGGATATCGGTTTGGGTCTTCGAGAAGACCTATTCC | Forward oligo containing BbsI stuffer |
| DD004.r | TCGAGGAATAGGTCTTCTCGAAGACCCAAACCGATATC | Reverse oligo containing BbsI stuffer |
| DD005.f | ATGAACGAAAAACAAAAGAGATTCG | Forward for amplifying *terS* from $\phi$NM1 for pDD1038 |
| DD005.r | TTAACTTTCGTCATCGTACTCAC | Reverse for amplifying *terS* from $\phi$NM1 for pDD1038 |
| DD006 | TATGAAGACACTGGAATGAACGAAAAACAAAAGAGATTCGC | Forward for amplifying *terS*/*terS*-*terL* from $\phi$NM1 ($P_{rpsL}$ overhang) |
| DD007 | TATGAAGACACTGTTAACTTTCGTCATCGTACTCACCAA | Reverse for amplifying *terS* from $\phi$NM1 for pDD1040 |
| DD008 | TATTGAAGACACGTTTAAGGATCTATGTGGGTTGGCT | Forward for amplifying $P_{terS}$-*terS* and $P_{terS}$-*terS*-*terL* from $\phi$NM1 |
| DD009 | TATTGAAGACACGAATTTAACTTTCGTCATCGTACTCACC | Reverse for amplifying $P_{terS}$-*terS* from $\phi$NM1 |
| DD0010 | TATGAAGACACGAGATGAACGAAAAACAAAAGAGATTCGC | Forward for amplifying *terS*-*terL* from $\phi$NM1 |
| DD0011 | TATGAAGACACGCTATAATCCTAGAGATTTTATTGTGTCAACTTTCGAACTGA | Reverse for amplifying *terS*-*terL* from $\phi$NM1 |
| DD0012 | TATTGAAGACACGAATCTATAATCCTAGAGATTTTATTGTGTCAACTTTCGAACTG | Reverse for amplifying $P_{terS}$-*terS*-*terL* from $\phi$NM1 |
| DD0013.f | AATATTGGTGAGTACGATGACGAAAGTTAAACGGTAACCAAGATGTCGAG | Forward for amplifying pDD1004 backbone for pDD1038 |
| DD0013.r | ATCTGCGAATCTCTTTTGTTTTTCGTTCATCTCGAGCATTCTAGACCATGG | Reverse for amplifying pDD1004 backbone for pDD1038 |

| | | |
|---|---|---|
| DD0014 | TATGAAGACACAACACGGTAACCAAGATGTCG | Forward for amplifying pDD1004 backbone for pDD1040 |
| DD0015 | TATGAAGACACTCCATGTGATATGTCCTCCTCTCT | Reverse for amplifying pDD1004 backbone for pDD1040/pDD2006 |
| DD0016 | TATGAAGACACTAGCACGGTAACCAAGATGTCG | Forward for amplifying pDD1004 backbone for pDD2005/pDD2006 |
| DD0017 | TATGAAGACACTCTCGAGCATTCTAGACCATGG | Reverse for amplifying pDD1004 backbone for pDD1040/pDD2006 |
| DD0018 | TATTGAAGACACGTTTAATTGACAGTAAAATGACAGTTTTTGACAC | Forward for amplifying $P_{terS\ \mathrm{full}}$-*terS* from $\phi$NM1 |
| DD0019.f | CTAGTCTAGAGAAGTGAAATAATGACACCGTG | Forward for amplifying 5' *terS* homology arm |
| DD0019.r | GTTTAATTTAACTTTCGTCATTCATTTCATTTACCACCAACTCTCGC | Reverse for amplifying 5' *terS* homology arm |
| DD0020.f | GTTGGTGGTAAATGAAATGAATGACGAAAGTTAAATTAAACTTTAACAAACC | Forward for amplifying 3' *terS* homology arm |
| DD0020.r | CCGCTCGAGAAACCCAATTCAGTTTAGATACTGG | Reverse for amplifying 3' *terS* homology arm |
| DD0021.f | GCAATCAAAGAGACATCAAAATATTCGG | Forward for amplifying fragment #1 from pDD0021 |
| DD0021.r | GCATCAGCCATGATGGATAC | Reverse for amplifying fragment #1 from pDD0021 |
| DD0022.f | GTATCCATCATGGCTGATGC | Forward for amplifying fragment #2 from pDD0021 |
| DD0022.r | CCGAATATTTTGATGTCTCTTTGATTGCCG | Reverse for amplifying fragment #2 from pDD0021 |
| DD0023.f | CTTTTTTTGAGATCTGTCCATACCCATGGTCTAGAAAGGATCTATGTGGGTTGGCTGAT | Forward for amplifying $P_{terS}$-*terS* for insertion into pDD0029 |
| DD0023.r | TGTTTGCTTTTGAAAAAAGATACAGGTATATTTTTCTGACTCGAGTTAACTTTCGTCATCGTACTCACCAATATTAATCT | Reverse for amplifying $P_{terS}$-*terS* for insertion into pDD0029 |
| DD0024.f | CTTTGTTATCTTGGTTACCCGTCT | Forward for amplifying pDD0021 backbone for *ori* swap |
| DD0024.r | GGTATTTACCACAACAGTACGCCA | Reverse for amplifying pDD0021 backbone for *ori* swap |
| DD0025.f | GGTTGGCGTACTGTTGTGGTAAATACCCCTTTGCGAAAGAGTTAATAAGTTAACAGAAG | Forward for amplifying pCN56 *ori* |
| DD0025.r | GCATATTAAGAAGACGGGTAACCAAGATAACAAAGCCAATAAAAGCAATCAATGAACCAAGAC | Reverse for amplifying pCN56 *ori* |
| DD0026.f | GCGAGAGTTGGTGGTAAATG | Forward for colony PCR check amplicon for *terS* deletion |

| | | |
|---|---|---|
| DD0026.r | TTAACTTTCGTCATCGTACTCACC | Reverse for colony PCR check amplicon for *terS* deletion |
| DD0027.f | GCGGTTTGTTATGACATACAC | Forward for *terS*-flanking check amplicon |
| DD0027.r | GGTAATCCATGTGATGTTCAATG | Reverse for *terS*-flanking check amplicon |
| DD0028.f | CGCCTGTACAACCATTTGGC | Forward for qPCR probe of *nuc* gene |
| DD0028.r | GCAAGTCCCTTTTCCACTAATTCC | Reverse for qPCR probe of *nuc* gene |
| DD0029.f | GCACAAATAGCGTCGGATGGG | Forward for qPCR probe of *SpCas9* gene |
| DD0029.r | AGCTGTCCGTTTGAGACGAG | Reverse for qPCR probe of *SpCas9* gene |
| SA_terS.f | GAAATATATAATGAATGGATGTAA | Forward primer for *terS* gRNA |
| SA_terS.r | AAACTTACATCCATTCATTATATA | Reverse primer for *terS* gRNA |
| SA_active.f | GAAATATCTTACTGCTGTTTTTTT | Forward primer for active gRNA |
| SA_active.r | AAACAAAAAAACAGCAGTAAGATA | Reverse primer for active gRNA |
| SA_null.f | GAAATGCTTGCCATAACATATCGA | Forward primer for null gRNA |
| SA_null.r | AAACTCGATATGTTATGGCAAGCA | Reverse primer for null gRNA |
